# 7-Tesla evidence for columnar and rostral–caudal organization of the human periaqueductal gray response in the absence of threat: a working memory study

**DOI:** 10.1101/2022.12.21.521479

**Authors:** Alexandra K. Fischbach, Ajay B. Satpute, Karen S. Quigley, Philip A. Kragel, Danlei Chen, Marta Bianciardi, Lawrence L. Wald, Tor D. Wager, Ji-Kyung Choi, Jiahe Zhang, Lisa Feldman Barrett, Jordan E. Theriault

**Affiliations:** Department of Psychology, Northeastern University, Boston, MA, USA; Department of Psychology, Emory University, Atlanta, GA, USA; Department of Radiology, Athinoula A. Martinos Center for Biomedical Imaging, Massachusetts General Hospital and Harvard Medical School, Boston, MA, USA; Department of Psychological and Brain Sciences, Dartmouth College, Hanover, NH, USA; Department of Surgery, University of California, San Francisco, CA, USA; Department of Psychiatry, Massachusetts General Hospital and Harvard Medical School, Boston, MA, USA; Athinoula A. Martinos Center for Biomedical Imaging, Massachusetts General Hospital and Harvard Medical School, Charlestown, MA, USA

## Abstract

The periaqueductal gray (PAG) is a small midbrain structure that surrounds the cerebral aqueduct, regulates brain–body communication, and is often studied for its role in “fight-or-flight” and “freezing” responses to threat. We used ultra-high field 7-Tesla fMRI to resolve the PAG in humans and distinguish it from the cerebral aqueduct, examining its *in vivo* function in humans during a working memory task (N = 87). Relative to baseline fixation, both mild and moderate task-elicited cognitive demands elicited bilateral BOLD increases in ventrolateral PAG (vlPAG), a region previously observed to show increased activity during anticipated painful threat in both non-human and human animals. The present task posed only the most minimal (if any) “threat”. The mild-demand condition involved a task easier than remembering a phone number, elicited a heart rate decrease relative to baseline, yet nonetheless elicited a bilateral vlPAG response. Across PAG voxels, BOLD signal intensity correlated with changes in physiological reactivity (relative to baseline) and showed some evidence of spatial organization along the rostral–caudal axis. These findings suggest that the PAG may have a broader role in coordinating brain—body communication during a minimally to moderately demanding task, even in the absence of threat.

## Introduction

Historically, the periaqueductal gray (PAG) has been identified as a key brain structure mediating the “fight-or-flight” and “freezing” responses (for reviews, see refs.^1, 2^), and this functional association of PAG with threat responses continues to be articulated in recent reviews^3–5^. Direct chemical stimulation of the PAG elicits a diverse range of motor behaviors and physiological responses—including, in cats, immobility, pupil dilation, vocalization (e.g., hissing or howling), alert positions^6^, running and jumping^7^, and hypertension and tachycardia^8^, all of which are generally understood as part of the “fight-or-flight” and “freezing” responses (for reviews, see refs.^9–11^). Likewise, in humans PAG is most commonly studied for its role in threat^12–18^. Deep-brain stimulation of the human PAG elicits a desire to escape^19^, feelings of fear and impending death, and autonomic changes including hyperventilation and increased heart rate^20^. Brain imaging studies in humans observed blood-oxygen level-dependent (BOLD) signal intensity increases in PAG when participants were chased through a virtual maze (i.e., approached by a simulated predator^21^), and when exposed to a learned cue signaling the onset of an unpleasant respiratory restriction (via an inspiratory resistance load^22, 23^). The interpretation of the PAG as a primary region for coordinating survival-based responses has been used to unify these many observations under a single coherent function^24, 25^, in which the PAG was conceived of as playing a central role in an evolutionarily ancient “fear circuit”^25–30^.

The hypothesis of PAG as a center for survival-based responses unifies a great deal of evidence, but not all evidence. For example, PAG has been observed to play a functional role in bodily processes at moments when threat is absent (for review, see ref.^31^), including control of the heart (e.g., heart rate^13, 32–34^), blood pressure control^35^, cardiorespiratory coordination^36^, bladder control^37, 38^, rapid-eye-movement (REM) sleep^39^, body temperature^40, 41^, trigeminovascular contributions to migraine headache^42^, communication (e.g., the vocalization of Songbirds^43^), mating behavior (e.g., the control of male copulatory behavior in Quails^44^), and feeding behaviors (e.g., the motivational drive to hunt in mice^45^). Furthermore, the PAG has extensive connectivity with regions across the entire neuraxis from spinal cord and brainstem to cerebral cortex, placing it at a critical integration point in brain–body communication (Table S1; for review, see ref.^46^). Building on this evidence, we hypothesize that evidence of PAG involvement in response to predatory threat is consistent with a more basic role of the PAG in coordinating and regulating internal bodily (visceromotor) systems in service of efficient regulation of the body (called allostasis) —particularly given that threat responses represent only one class of contexts in which the autonomic nervous system, the endocrine system, the immune system and tissues of the body must be coordinated with each other and with motor movements. We hypothesize that the PAG plays a more general regulatory role that extends beyond extreme or threatening survival-related circumstances^47, 48^ (for reviews, see refs.^49, 50^) and tested this hypothesis by examining the BOLD response in functionally-specific PAG columnar subregions during a mild-to-moderately challenging working memory task.

During survival-based investigations, the PAG is routinely subdivided into functionally specialized columnar subregions (see Fig. 2). Research has identified dorsomedial (dmPAG), dorsolateral (dlPAG), lateral (lPAG), and ventrolateral (vlPAG) columns (for reviews, see refs.^2, 31^), such that dl/lPAG is thought to support active coping “fight-or-flight” responses to predatory threats, dmPAG is thought to support similar responses to aggressive conspecifics, and vlPAG is thought to support passive coping or “freezing” responses to painful threats^7, 51, 52^. For example, stimulation of dl/lPAG, in rodents and cats increases cardiac and respiratory rates^53–57^ and elicits explosive running and jumping^7, 52, 53, 58^. Stimulation of vlPAG in rodents and cats decreases cardiac and respiratory rates^53–56, 59^, and induces an absence of movement in cats^7^ and “freezing” behavior in rats^52, 60^. In humans, the increase in PAG BOLD signal during anticipated breathlessness is also localized to vlPAG^22, 23^. Interestingly, subcutaneous (deep muscle) and cutaneous pain also elicit a human PAG BOLD response in vlPAG and lPAG respectively^61^.

In the present study, we examined changes in PAG columnar activity in the context of a mild-to-moderately challenging *N*-back working memory task, which aims to simulate energetic, cognitive, and physiological demands within the range of those typically encountered in daily life. In this task participants viewed a sequence of letters presented one at a time and responded with a key press when a letter was repeated either from the immediately prior trial (1-back) or three trials prior (3-back; Fig. 1). This task required that participants simultaneously attend to and remember target stimuli while inhibiting or completing task-relevant motor responses. Given that the 1-back task could be conceptually described as easier than remembering a phone number, we consider it to fall well outside the traditional survival-based contexts the PAG function has previously been probed within.

**Figure 1.**
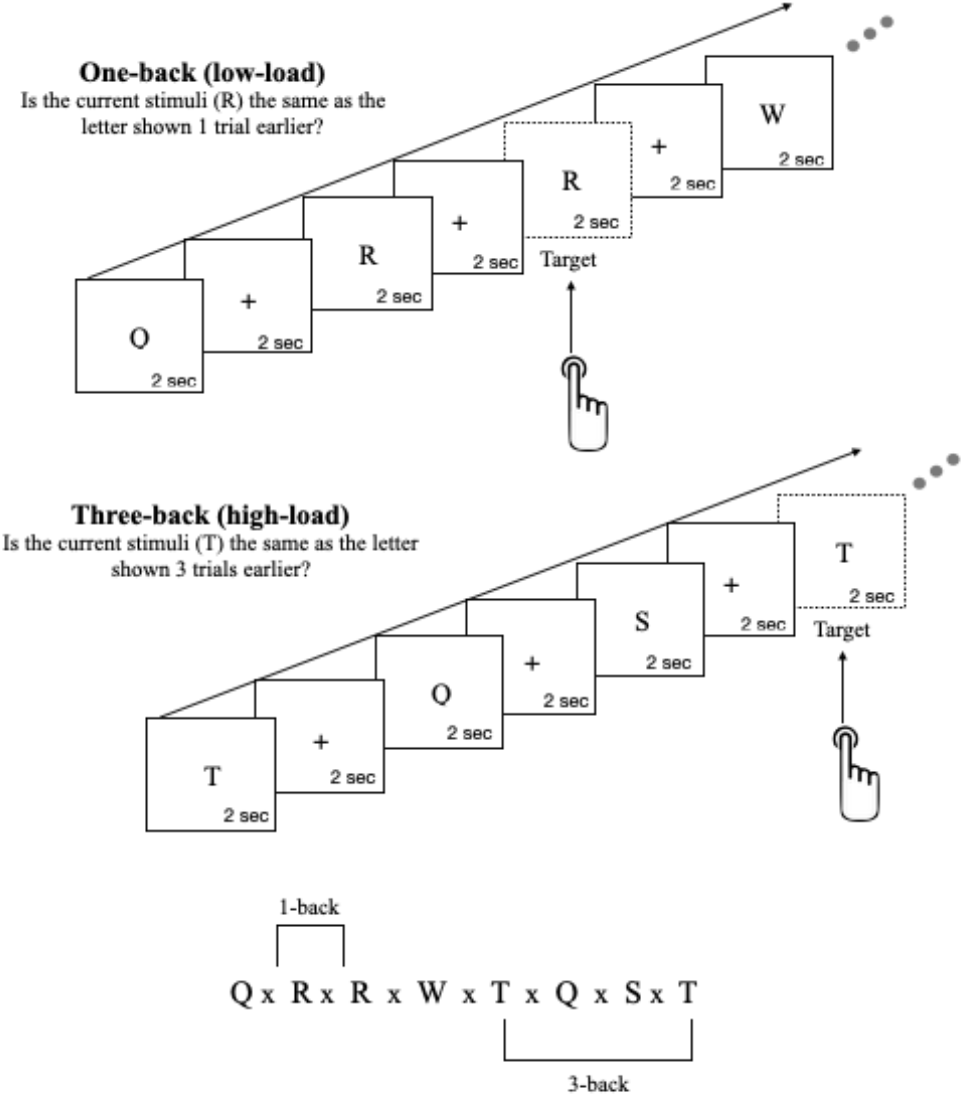
Schematic overview of the working memory *N*-back task. An example of a 1-back (mild cognitive demand) and 3-back (moderate cognitive demand) task showing a sequence of trials. For every trial, a single letter was presented at the center of the visual field for 2-seconds followed by a fixation cross for 2-seconds. Participants respond with a button press if the target stimulus matched the stimulus *n* (either 1 or 3) positions back.

Imaging using 1.5 Tesla and 3 Tesla MRI scanners struggles to resolve small brainstem nuclei, both due to low signal-to-noise ratios (SNR) and voxel resolution, whereas the ultra-high resolution 7 Tesla (7T) fMRI used in the present work provides a substantial increase in SNR^62, 63^. An earlier report included the first 24 participants from the current data set (all the participants who were available for analysis at that time) and used univariate and multivariate methods to distinguish PAG BOLD responses to 1-back (mild cognitive demand) and 3-back (moderate cognitive demand) conditions^64^. We reported initial evidence that BOLD signal intensity increased for the 3-back vs. the 1-back condition and that this PAG response was localized to the right rostral vlPAG. The present work builds on this earlier report, with a sample size (N = 87) that affords the power to identify functional distinctions among PAG columns. We also improved our localization procedure, identifying PAG columnar subregions by their radial degree. As in prior work^65^, we identified subject-specific PAG masks and aligned them at the group level to estimate the voxel-wise BOLD response during 1-back and 3-back conditions of the *N*-back working memory task. For PAG columns, tract-tracing techniques in non-human animals typically produce symmetrical results^7, 54, 66–71^, and given this we expected that a well-powered analysis of the PAG response to mild and moderate working memory load would produce a bilateral symmetric response as well, if the task elicits a response from functional columns of the PAG. Following earlier work^65^, we also recorded cardiac interbeat interval (IBI) and respiratory rate (RR) to examine whether columnar organization supported visceromotor functions, and whether this visceromotor support explained any task-elicited changes observed during the working memory task.

## Results

### Behavior significantly differed between 1-back and 3-back working memory conditions

Consistent with the earlier work using a small subset of this sample^64^, behavioral results differed between 1-back and 3-back task conditions such that participants gave more correct answers (i.e., higher hit rates) for 1-back trials (*M* = 0.992, *SD* = 0.022), than for 3-back trials (*M* = 0.892, *SD* = 0.080; *t_(85)_* = 12.18, *p* < 1e^−16^, *d* = 1.31). Participants also responded more quickly for correct 1-back (*M* = 0.762 sec, *SD* = 0.156 sec) than for correct 3-back trials (*M* = 0.993 sec, *SD* = 0.215 sec; *t_(85)_* = 13.75, *p* < 1e^−16^, *d* = 1.48). Response times for incorrect responses did not significantly differ between 1-back and 3-back conditions.

### Peripheral physiological responses (relative to baseline) were observed during both the 1-back and 3-back working memory conditions

Peripheral physiological changes in IBI and RR during the task (relative to baseline) were computed for each participant in 1-back and 3-back task epochs (see Methods) and tested against a null hypothesis of zero change from the baseline. Unlike the preliminary findings^65^ which observed no changes in IBI and RR relative to baseline, we observed physiological changes in both working memory conditions (see Fig. 3). On average across participants, the 1-back condition was associated with cardio-deceleration and increased respiratory rate (IBI: *t*_(58)_ = 3.637, *p* < 0.0006, *d* = 0.47; RR: *t*_(46)_ = 4.750, *p* < 1e^−5^, *d* = 0.69) whereas the 3-back condition was associated with cardio-acceleration and increased respiratory rate (IBI: *t*_(58)_ = 3.825, *p* < 0.0003, *d* = 0.50; RR: *t*_(46)_ = 9.067, *p* < 1e^−12^, *d* = 1.32). Furthermore, the 3-back condition produced a greater reduction in IBI (*M* = −12.5 ms, *SD* = 25.0 ms; i.e., faster HR relative to baseline) when compared to the 1-back condition (*M* = 9.64 ms, *SD* = 20.4 ms; *t*_(58)_ = 5.21, *p* < 1e^−6^, *d* = 0.68; Fig. 3). Respiration rate was also relatively faster in the 3-back condition (relative to baseline; *M* =1.94 breaths per minute (bpm), *SD* = 1.47 bpm) compared to the 1-back condition (*M* = 0.90 bpm, *SD* = 1.30 bpm; *t*_(46)_ = 6.07, *p* < 1e^−7^, *d* = 0.89; Fig. 3).

**Figure 2.**
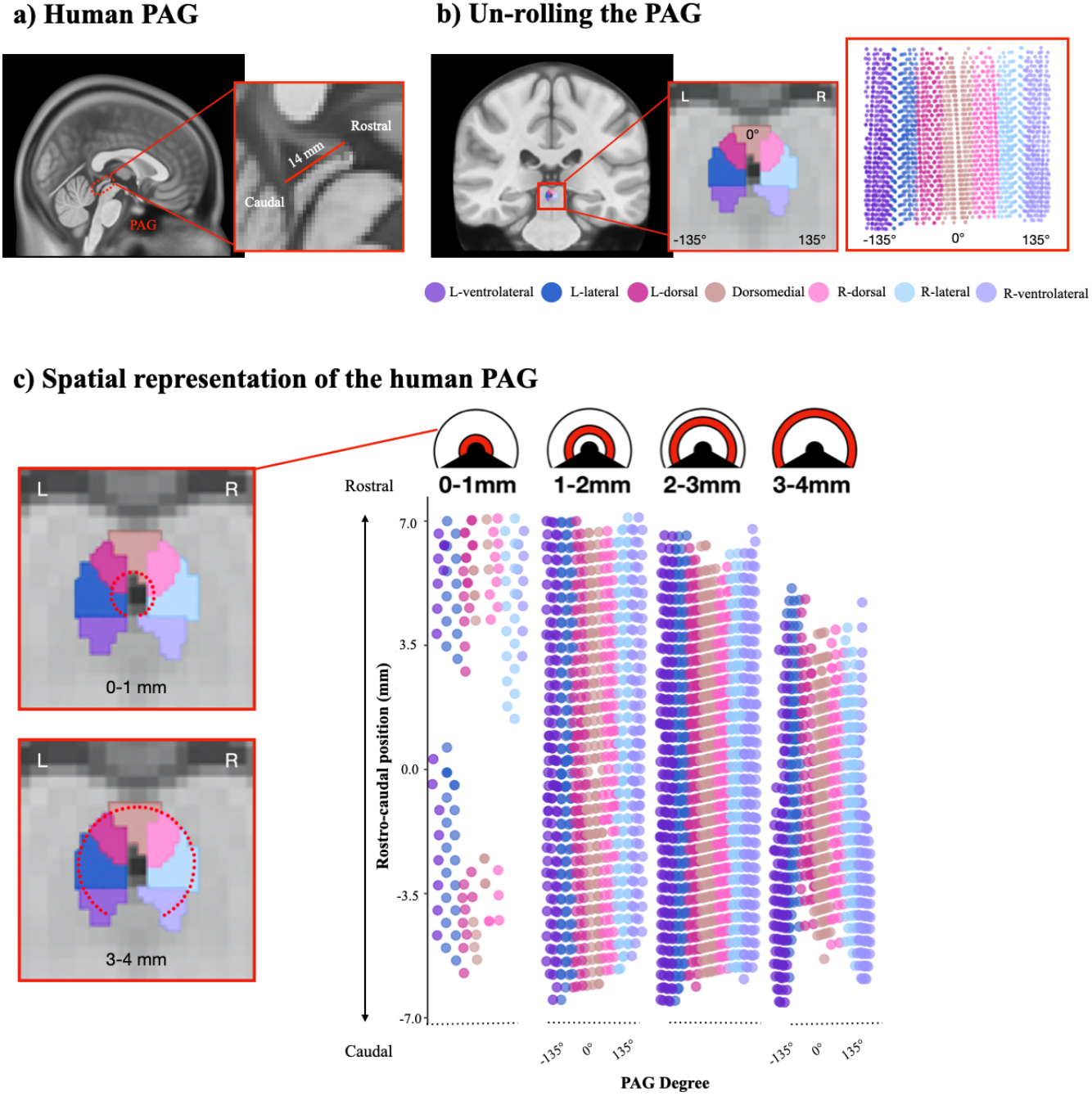
PAG identification, segmentation, and visualization. (**a**) 7-Tesla fMRI sagittal view of the human PAG. (**b**) Group-level template of the human PAG (coronal view). The PAG template can be divided radially into 7 columnar subregions (e.g., L-vlPAG, L-lPAG, L-dPAG, dmPAG, R-dPAG, R-lPAG, R-vlPAG). (**c**) 2-D visualization of PAG voxels. Upper panels display radial cross-sections of the PAG ‘cylinder’ (e.g., the 0-1 mm panel shows voxels abutting the cerebral aqueduct, and the 1-2 mm panel shows voxels 1-2 mm from the center of the cerebral aqueduct). The y-axis illustrates the longitudinal (rostral–caudal) axis of the PAG. The x-axis represents voxel position in +/− 135 degrees from the brain midline (ventrolateral = +/− 97.5-135°; lateral = +/− 60-97.5°; dorsolateral = +/− 22.5-60°; dorsomedial = +/− 22.5°). Abbreviations: PAG: periaqueductal gray.

**Figure 3.**
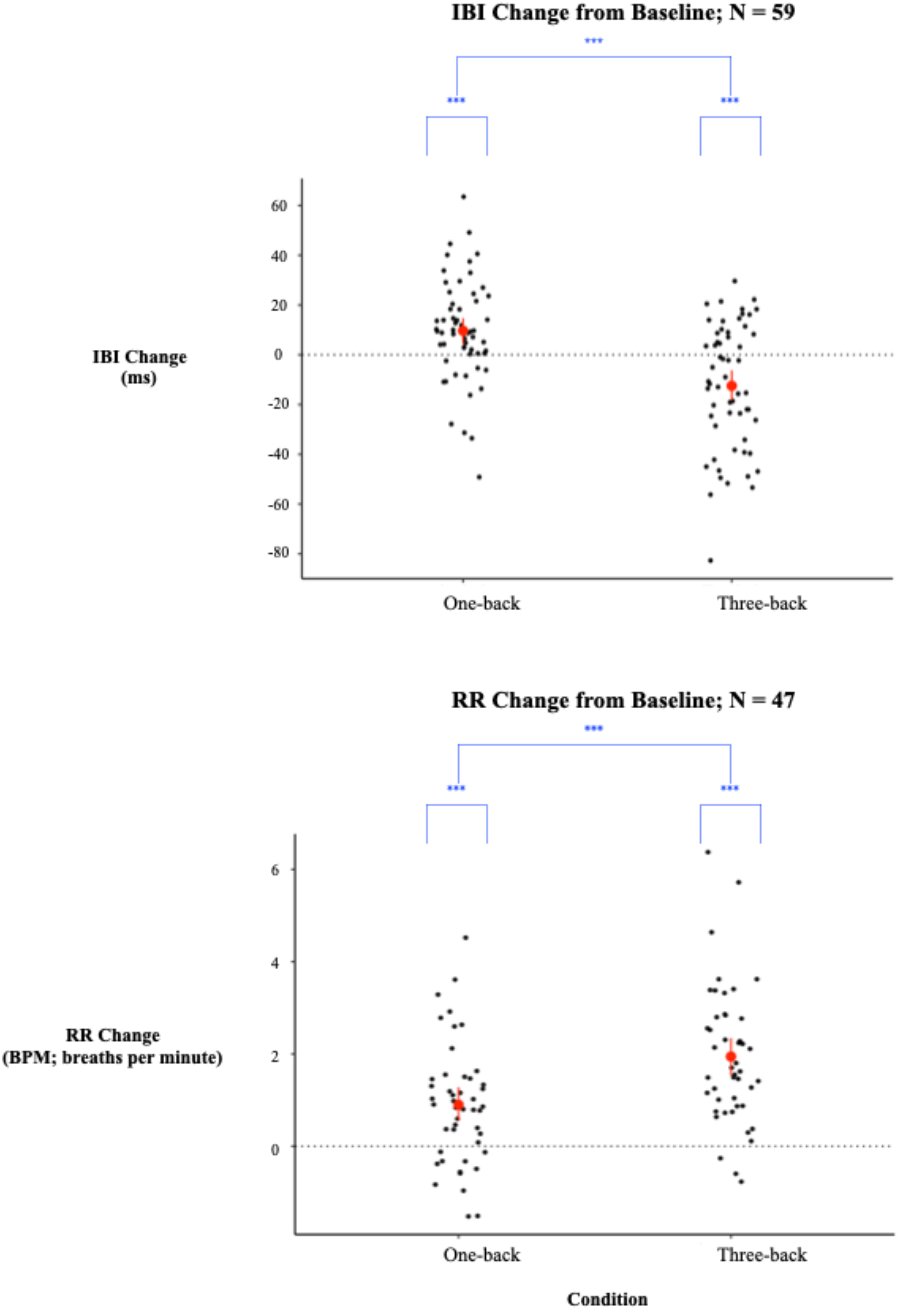
Task-dependent physiological changes from baseline. Mean reactivity (with a 95% confidence interval) for IBI and RR data (see *Peripheral Physiological Recording & Quality Assessment*) measured during the 1-back (left) and 3-back (right) conditions of the *N*-back task. Consistent with our chosen analytic approach (see *Peripheral Physiological Preprocessing and Analysis)*, cardiac data are presented as measurements of IBI; an increase in IBI reflects a decrease in heart rate and a decrease in IBI reflects an increase in heart rate. Black dots illustrate individual mean change scores (for IBI or RR, where each measure is a change score from baseline in measure-specific units). Change scores are significant at *p* < 0.001***. Abbreviations: IBI: interbeat interval; RR: respiration rate.

### PAG BOLD signal intensity, compared to baseline fixation, increased during the 1-back and 3-back working memory conditions

Consistent with the preliminary findings reported in prior work^64^, average PAG BOLD signal intensity (across all PAG voxels) increased during the 3-back condition relative to baseline fixation (*t_(86)_* = 2.41, *p* < 0.01, *d* = 0.26). In contrast to this earlier work, PAG BOLD signal intensity also increased from baseline fixation in the 1-back condition (*t_(86)_* = 1.79, *p* < 0.039, *d* = 0.19), and of particular note, there was no significant difference in overall PAG BOLD signal intensity between 1-back and 3-back conditions (*t_(86)_* = 1.34, *p* < 0.091, *d* = 0.14). That is, both 1-back and 3-back working memory conditions increased the average PAG BOLD response, without evidence of significant overall differences between the two conditions.

### PAG BOLD signal intensity revealed columnar specificity

Expanding on preliminary findings^65^, visual inspection of BOLD signal intensity (vs. baseline fixation) in 1-back and 3-back task periods revealed what appeared to be columnar stripes of BOLD signal intensity increases that were strongest in bilateral vlPAG, and extended along the entire rostral–caudal axis (Fig. 4a). To quantify this observation (see Methods), we subdivided the PAG along its radial degrees (*degree rank;* comprised of 10 ordinal ranks) and along its rostral–caudal axis (*rostral–caudal rank;* comprised of 4 ordinal ranks; Fig. 4b). PAG BOLD signal intensity was averaged within subregions (combining left/right sides for this analysis) and submitted to a linear mixed-effects model including all main effects and interactions [*N*-back condition (3-back vs. 1-back) x degree rank (1-10) x rostral–caudal rank (1-4)].

**Figure 4.**
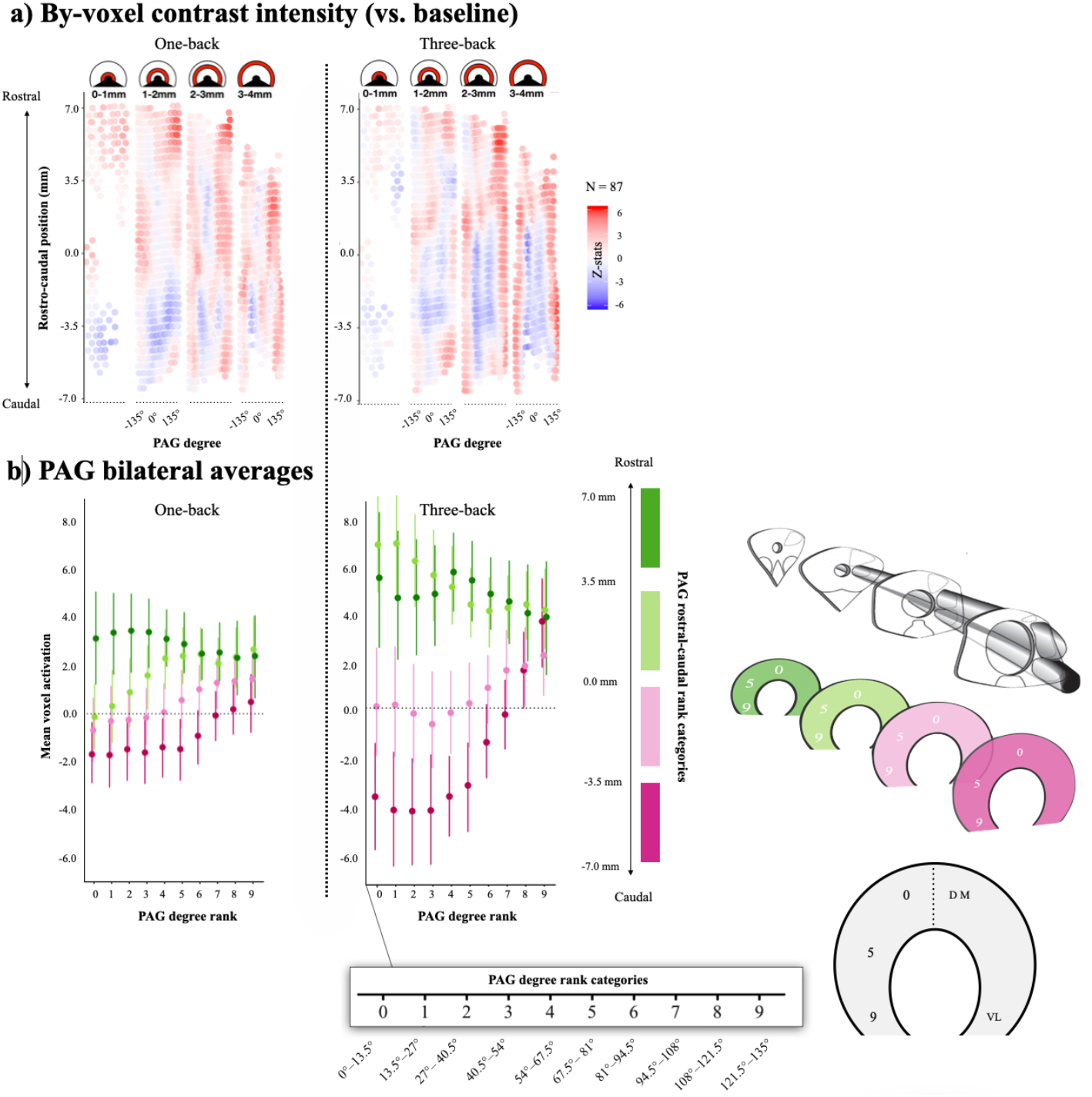
Task-elicited PAG BOLD signal intensity. (**a**) Voxelwise PAG BOLD signal intensity during 1-back (left) and 3-back (right) conditions in the *N*-back task. Upper panels display radial cross-sections of the PAG ‘cylinder’ (e.g., 0-1 mm). The y-axis illustrates voxel position along the rostral–caudal axis. The x-axis represents voxel position in +/− 135 degrees from the brain midline (ventrolateral = +/− 97.5-135°; lateral = +/− 60-97.5°; dorsolateral = +/− 22.5-60°; dorsomedial = +/− 22.5°). The heat map illustrates PAG BOLD signal intensity relative to baseline fixation between task blocks. (**b**) Mean voxelwise activation during 1-back (left) and 3-back (right) conditions, as modeled by PAG degree and rostral–caudal position.

We observed a significant three-way interaction [*F_(1, 86.00)_* = 5.43, *p* < 0.022]. Within the 3-back condition, we observed significant main effects of degree rank [*F_(1,86.00)_* = 17.09, *p* < 1e^−5^] and of rostral–caudal rank [*F_(1,86.00)_* = 24.03, *p* < 1e^−6^], qualified by an interaction [*F_(1, 86.00)_* = 14.99, *p* < 0.0002]. Relative to dmPAG (i.e., degree rank of 0), BOLD signal intensity was higher in more lateral (i.e., higher degree rank) PAG regions [simple slope of degree rank; *B* = 0.64, *SE* = 0.16, *t_(86.00)_* = 4.13, *p* < 1e^−5^], and this effect of degree rank was higher at the caudal end of the PAG (degree rank x rostral–caudal interaction; for a visualization of this relationship, see Fig. 4b, middle panel). BOLD signal intensity was also higher toward the rostral end of the PAG [simple slope of rostral–caudal rank; *B* = 3.86, *SE* = 0.79, *t_(86.00)_* = 4.90, *p* < 1e^−6^].

Within the 1-back condition, we also observed significant main effects of degree rank [*F_(1,86.00)_* = 9.27, *p* < 0.003] and of rostral–caudal rank [*F_(1,86.00)_* = 12.14, *p* < 0.0008]. In the 1-back condition, the interaction between degree rank and rostral–caudal rank was marginally significant [*F_(1,86.00)_* = 3.60, *p* < 0.061]. Similar to the 3-back condition, the 1-back condition elicited higher BOLD signal in more lateral (i.e., higher degree rank) PAG regions relative to dmPAG [simple slope of rostral–caudal rank: *B* = 0.34, *SE* = 0.11, *t_(86.00)_* = 3.04, *p* < 0.003], and this effect of degree rank was higher at the caudal end of the PAG (degree rank x rostral–caudal interaction; Fig. 4b, left panel). Also as in the 3-back condition, the 1-back condition elicited higher BOLD signal intensity toward the rostral end of the PAG [*B* = 1.81, *SE* = 0.52, *t_(86.00)_* = 3.485, *p* < 0.0007].

### Peripheral physiological responses (relative to baseline) correlated to PAG BOLD signal intensity along medial and caudal sections of the rostral–caudal axis

Given the role of the PAG in physiological regulation, we examined whether the task-elicited visceromotor changes we observed were associated with task-elicited changes in PAG bold signal during the working memory task. In both conditions (relative to baseline), we examined the voxel-wise spatial organization of correlations between PAG BOLD signal intensity and peripheral physiology (i.e., cardiac interbeat interval (IBI) and respiratory rate (RR)). Correlations were thresholded using a bootstrapped confidence interval, resampling across participants [1000 resamples], and excluding any voxel where the 95% confidence interval included zero. We observed some evidence for spatially-organized correlations along the rostral–caudal axis of the PAG, but unlike the task-dependent PAG BOLD signal, IBI and RR changes correlated with PAG BOLD signal intensity across multiple columns (Fig. 5; shaded overlay). Increases in IBI were correlated with increased PAG BOLD activity in the more medial and more caudal areas across multiple columns during the 3-back condition (Fig. 5, upper right panel). Comparatively, in the 1-back task period, there were fewer statistically significant correlations observed (Fig. 5, upper left panel). Change in RR (relative to baseline) also showed some evidence of spatially-organized correlations along the rostral–caudal axis of the PAG, with scattered positive correlations in medial and caudal areas of the rostral–caudal axis during both the 3-back and 1-back conditions (Fig. 5, bottom panel).

**Figure 5.**
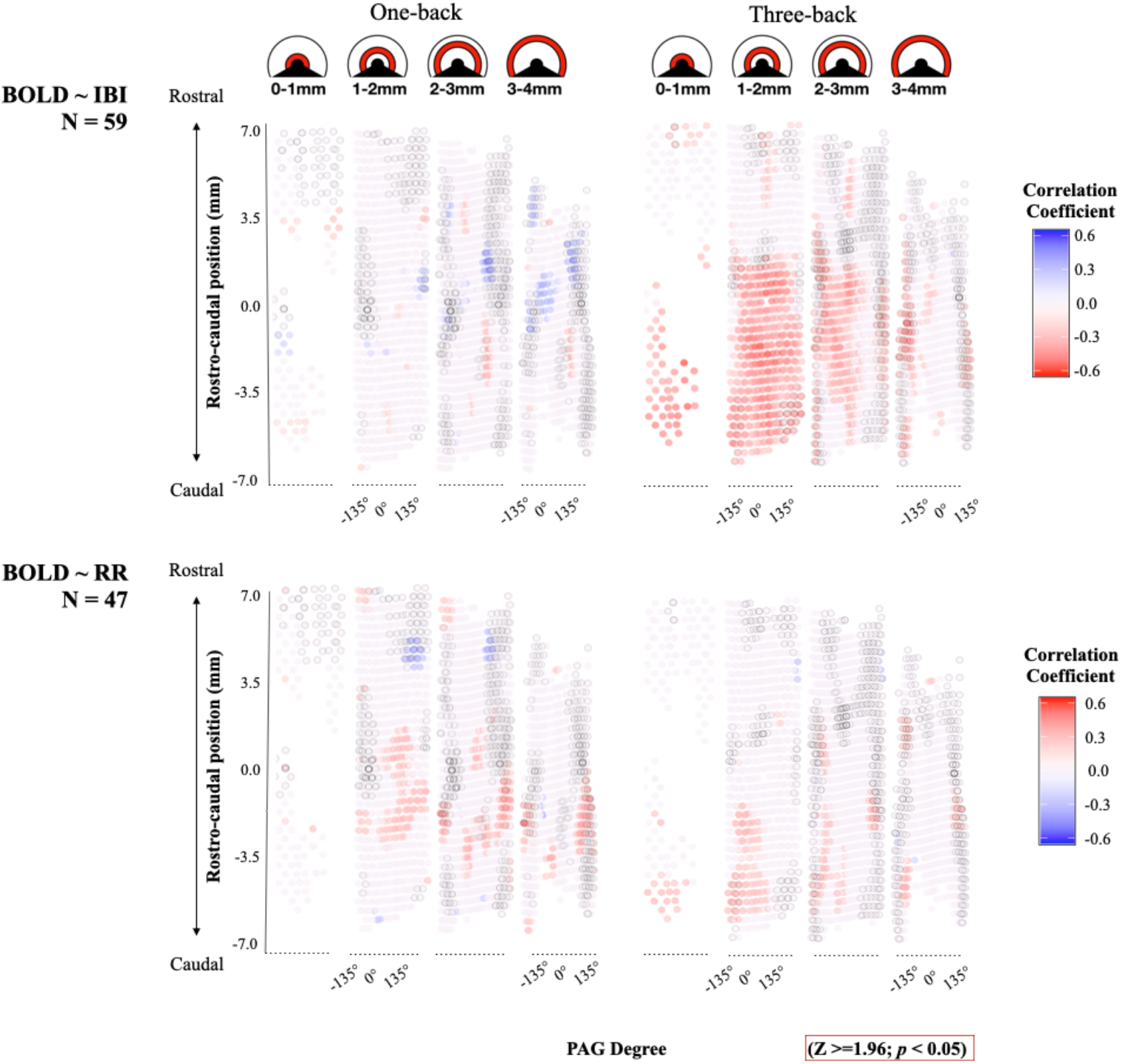
Between-subjects bootstrapped correlations between peripheral physiology voxelwise PAG BOLD signal intensity. Upper panels display radial cross-sections of the PAG ‘cylinder’ (e.g., 0-1 mm). The y-axis illustrates voxel position along the rostral–caudal axis. The x-axis represents voxel position in +/− 135 degrees from the brain midline (ventrolateral = +/− 97.5-135°; lateral = +/− 60-97.5°; dorsolateral = +/− 22.5-60°; dorsomedial = +/− 22.5°). The heat map illustrates magnitude (i.e., strength) of the correlation coefficients. Correlation coefficients are thresholded using bootstrap resampling across subjects within each voxel (1000 resamples). Coefficients that include 0 in their 95% confidence interval were masked out. Voxels from the 1-back/3-back contrast heat map (Figure 4a) were thresholded at (Z >=1.96; *p* < 0.05) and superimposed onto the current figure; the voxels that met the thresholded criterion are illustrated by black circles. Notably, the correlation coefficients for IBI and RR do not do not exhibit the distinct columnar spatial organization of bilateral vlPAG as seen in task-elicited PAG BOLD (Figure 4a). Abbreviations: IBI: interbeat interval; RR: respiration rate.

## Discussion

Our results suggest that the human PAG is involved during tasks that evoke modest changes in physiological activity and that are neither painful nor threatening. We observed greater BOLD signal changes, most notably in vlPAG, during both the moderately demanding 3-back condition and the minimally demanding 1-back condition of an *N*-back working memory task. The BOLD signal increases during both mild and moderate cognitive demand were most notable in vlPAG, suggesting some columnar specificity of these effects. These findings are consistent with evidence that PAG neurons contribute to coordinating and regulating the internal (visceromotor) systems of the body, including the autonomic nervous system (for reviews, see refs.^22, 72, 73^). In other words, although PAG circuitry may be intensely engaged during threatening episodes involving “fight-or-flight” or “freezing”, the function of the PAG may not be specific to those behaviors.

In both *N*-back conditions, BOLD signal increases were observed bilaterally in the vlPAG (Fig. 4a), with the greatest increases at the caudal pole (Fig. 4b). This spatially-organized pattern of PAG BOLD signal intensity changes is consistent with an earlier 7T scanning study from our group, where an increase in ventrolateral (vlPAG) BOLD signal at the caudal pole was observed when participants viewed affectively evocative images^63^. Indeed, the spatial organization of PAG BOLD signal intensity observed in prior work^63^ and the current study is also consistent with anatomical observations in non-human animals. For instance, in rodents, immunohistochemistry is used to detect cells positive for Fos, an immediate early gene used in signal transduction often used as a marker of recent local neuronal activity in brain tissues^74^. In rats, the distribution of Fos-positive cells in the PAG after administering a mild stressor (e.g., foot shock) resembled the spatial organization we observed during the working memory task, with the vlPAG exhibiting the highest concentration of Fos-positive cells, particularly in its caudal pole^60^. Taken together, our observation of spatially-organized BOLD signal intensity increases with concentrations in the caudal vlPAG during a working memory task, is consistent with and builds upon previous anatomical and functional observations in both human and non-human PAG.

Both *N*-back conditions elicited physiological changes in the current study. Both mild (1-back) and moderately demanding (3-back) conditions elicited increases in RR from baseline, whereas changes in IBI from baseline differed between conditions, with IBI increases in the 1-back (cardiodeceleration) and decreases in the 3-back (cardioacceleration). During the 3-back condition, both IBI and RR changes were more strongly correlated with increased PAG BOLD signal in medial and caudal regions of the PAG, but statistically significant correlations were observed relatively more frequently for changes in IBI than for changes in RR. The relative focus of BOLD signal increase in the vlPAG during 1-back and 3-back conditions was absent from the voxel-wise PAG correlations with physiological measures (Fig. 5; shaded overlay). This suggests that the BOLD response in vlPAG observed for 1-back and 3-back conditions is not purely a product of task-elicited cardiac or respiratory changes.

Within the past decade, significant progress has been made in imaging the human PAG (e.g., refs.^12–18, 61^); however, a critical methodological limitation is that these studies often use experimental contexts involving various threatening stimuli to induce PAG BOLD signal changes. By contrast, the *N*-back working memory task involves cognitive demands that are within the range of what is experienced in daily life, allowing us to observe the PAG BOLD response during less affectively evocative task conditions. Successful performance in a *N*-back working memory task requires integrating exteroceptive (e.g., visual stimuli) and interoceptive sensory signals (e.g., signals of energy status from the body), maintaining those signals across trials (i.e., working memory), mobilizing energy to meet changing task demands (e.g., as reflected in changes in cardiorespiratory reactivity^36, 75–77^), maintaining goal-directed behavior (e.g., allocating attentional resources), and finally, generating and executing a motor plan (e.g., pushing a button to make a response). These same brain functions not only support successful working memory performance, but they also support many other complex, goal-directed sensorimotor behaviors. Indeed, these are the basic operations that are needed to perform many laboratory psychological tasks and engage in real-world motivated performance (i.e., situations that are goal-relevant to the performer, require instrumental cognitive responses, and are active rather than passive^78^). From this perspective, the working memory task is a motivated performance task, similar to other motivated performance tasks that have elicited a PAG BOLD response under more stressful conditions (e.g., social stress during preparation for public speaking^34, 79, 80^). The *N*-back task, then, in addition to being regarded as a prototypic ‘cognitive’ task, can also be viewed in a broader context as a task that requires a brain to mobilize resources to meet task demands, which requires allostatic efforts^81, 82^. The PAG is anatomically well-positioned to contribute to allostasis —especially given the PAG’s location as an important convergence point for ascending viscerosensory signals from the body and descending visceromotor signals for controlling the body (for review, see ref.^46^).

Taken together, our observation of PAG BOLD response during *both mild and moderately challenging* cognitive tasks is consistent with a view of the PAG as a vital integrative structure that integrates sensory signals from the body with visceromotor control signals to the body in support of complex coordinated behaviors (for reviews, see refs.^49, 83–85^). Future research should prioritize the exploration of PAG function in naturalistic conditions, sampling situations that vary in resource demand and mobilization (as in the present work). Data from such experimental contexts would help characterize subcortical responses that are commonly missed in laboratory conditions that probe or simulate only the most extreme survival-relevant contexts.

## Acknowledgements

This work was supported by grants from the National Science Foundation (BCS 1947972), the National Institutes of Health (R01 AG071173, R01 MH109464, and R01 MH113234), the U.S. Army Research Institute for the Behavioral and Social Sciences (W911NF-16-1-019), and the Unlikely Collaborators Foundation. The views, opinions, and/or findings contained in this manuscript are those of the authors and shall not be construed as an official Department of the Army position, policy, or decision, unless so designated by other documents, nor do they necessarily reflect the views of the Unlikely Collaborators Foundation

## Methods

### Participants

One hundred forty participants were recruited from the Greater Boston area and provided informed consent in accordance with guidelines set forth by the Partners’ Healthcare Institutional Review Board (IRB). Consented participants completed two scanning sessions and were paid $150 for each scan (i.e., $300 upon study completion). Eighteen participants either withdrew or ended the MRI scan session before starting the working memory task. Nineteen participants were excluded due to poor image quality (e.g., large artifacts compromising analysis), as assessed by visual inspection in Mango version 4.1 (RRID: SCR_009603). Sixteen participants were excluded for excessive head motion (> 0.5 mm framewise displacement in > 20% of TRs in the run; see *MRI preprocessing*). Our final sample consisted of 87 participants (*M_age_* = 27.0 years, *SD_age_* = 6.1 years; 39 female, 47 male, 1 non-response). For self-reported ethnicity, 13% of participants identified as Hispanic. For self-reported race, 62% identified as White, 25% identified as Asian, and 9% identified as Black. Of the 87 participants, three did not provide a response for race, and one did not provide a response for ethnicity. The participant sample was generally well-educated (24% had some graduate education, 25% had completed college/university, 37% had some college/university, 10% had completed high school, and 2% had not completed high school). Peripheral physiological recordings were inspected for artifacts (see *Peripheral Physiological Recording and Quality Assessment*) and only non-artifactual recordings were included in analyses. Of the 87 participants with usable fMRI data, 59 had artifact-free cardiac data, and 47 had artifact-free respiratory data.

All participants were between 18 and 40 years old, right-handed, fluent English speakers, with normal or corrected to normal vision, and no known neurological or psychiatric illnesses. Participants were excluded from participating if they were pregnant, claustrophobic or had any metal implants that could cause harm during scanning.

### Experimental Paradigm

Participants completed a visual *N*-back working memory task during fMRI scanning, based on the experimental design of prior studies^86–90^ (Fig. 1). On every trial, a letter (Q, W, R, S, or T) was presented at the center of the visual field for 2-seconds followed by a fixation cross for 2-seconds. Participants were instructed to respond with a button press when the letter on the screen matched the one presented *n* trials ago where *n* was either 1 or 3. The task was administered during a single scanning session, consisting of a total of 12 blocks. Each block began with a cue indicating the current trial type (1-back or 3-back) and each block consisted of 10 trials (totaling 120 trials upon completion of the task). 1-back and 3-back blocks were presented in ABBA or BAAB order (counterbalanced across participants) and each block was followed by 25-seconds of resting fixation. Stimuli were classified into 2 categories; a “target” was a stimulus that met the *N*-back criteria for a match (e.g., *N*-back either 1 or 3 trials ago), and all other stimuli were classified as “nontargets”. The task had fixed proportions of 20% targets and 80% nontargets. Of the nontargets, 12.5% were classified as lure trials; lure trials were defined as 2-back matches in the 1-back blocks and lures in 3-back blocks were either 2 or 4-back matches. The proportion of lure trials was the same for both 1-back and 3-back blocks.

The task was administered in MATLAB (The MathWorks; RRID:SCR_001622), using the Psychophysics Toolbox extensions (ref.^91^; RRID:SCR_002881). Stimuli were projected such that participants viewed them on a mirror affixed to the head coil used for data acquisition. Response times and hit rates (calculated as the number of correct responses divided by the number of targets) were recorded using an MR-compatible button box. Before scanning onset to ensure familiarity with the task, participants completed a practice *N*-back task session on a laptop. These practice sessions mirrored the task used during scanning, including blocks of 1-back and 3-back trials, for a total of 48 trials.

### MRI Data Acquisition

Gradient-echo echo-planar imaging BOLD-fMRI was performed on a 7-Tesla (7T) Siemens MRI scanner at the Athinoula A. Martinos Center for Biomedical Imaging at Massachusetts General Hospital (MGH), Boston, MA. The scanner was built by Magnex Scientific (Oxford, UK), with the MRI console, gradient and gradient drivers, and patient table provided by Siemens. Functional images were acquired using GRAPPA-EPI sequence: echo time = 28 ms, repetition time = 2.34 s, flip angle = 75°, slice orientation = transversal (axial), anterior to posterior phase encoding, voxel size = 1.1 mm isotropic, gap between slices = 0 mm, number of axial slices = 123, field of view = 205 × 205 mm^2^, GRAPPA acceleration factor = 3; echo spacing = 0.82 ms, bandwidth = 1414 Hz per pixel, partial Fourier in the phase encode direction = 7/8. A custom-built 32-channel radiofrequency coil head array was used for reception. Radiofrequency transmission was provided by a detunable band-pass birdcage coil.

Structural images were acquired using a T1-weighted EPI sequence, selected so that functional and structural images had similar spatial distortions, which facilitated co-registration and subsequent normalization of data to MNI space. Structural scan parameters were: echo time = 22 ms, repetition time = 8.52 s, flip angle = 90°, slice orientation = transversal (axial), voxel size = 1.1 mm isotropic, gap between slices = 0 mm, number of axial slices = 126, field of view = 205 × 205 mm^2^, GRAPPA acceleration factor = 3; echo spacing = 0.82 ms, bandwidth =1414 Hz per pixel, partial Fourier in the phase encode direction = 6/8.

### MRI Preprocessing

Functional and structural MRI data were preprocessed using the fmriprep pipeline, version 1.1.2 (refs.^92, 93^; RRID: SCR_016216), a Nipype-based tool (refs.^93, 94^; RRID: SCR_002502). Full pipeline details can be found at (https://fmriprep.org/en/1.1.2/workflows.html). Spatial normalization of anatomical data to the 2009c ICBM 152 Nonlinear Asymmetrical template (ref.^95^; RRID: SCR_008796) was performed through nonlinear registration with the `antsRegistration` tool of ANTs version 2.1.0 (ref.^96^; RRID: SCR_004757), using brain-extracted versions of both T1w (T1-weighted) volume and template. Brain tissue segmentation of cerebrospinal fluid (CSF), white-matter (WM) and gray-matter (GM) was performed on the brain-extracted T1w using `FAST` in FSL version 5.0.9 (ref.^97^; RRID: SCR_002823). Functional data were slice time corrected using `3dTshift` from AFNI version 16.2.07 (ref.^98^; RRID: SCR_005927) and motion corrected using `MCFLIRT` (FSL version 5.0.9^99^). This was followed by co-registration to the corresponding T1w using boundary-based registration^100^ with 9 degrees of freedom, using `FLIRT` (FMRIB’s Linear Image Registration Tool; FSL version 5.0.9^99, 101^). Motion correcting transformations, BOLD-to-T1w transformation, and T1w-to-template (MNI) warp were concatenated and applied in a single step using `antsApplyTransforms` (ANTs version 2.1.0) using Lanczos interpolation. Physiological noise regressors were extracted using the aCompCor method^102^, taking the top five principle components from subject-specific CSF and WM masks, where the masks were generated by thresholding the WM/CSF masks derived from `FAST` at 99% probability, constraining the CSF mask to the ventricles (using the ALVIN mask^103^), and eroding the WM mask using the `binary_erosion` function in SciPy version 1.6.^104^. Frame-wise displacement^106^ was calculated for each functional run using the implementation of Nipype. Many internal operations of fmriprep use Nilearn (ref.^107^; RRID: SCR_001362), principally within the BOLD-processing workflow. For all participants, the quality of brain masks, tissue segmentation, and MNI registration was visually inspected for errors using the html figures provided by the fmriprep pipeline.

### PAG Identification and Alignment

The current approach was founded on manual methodologies developed in a previous 7 Tesla imaging study (for details, see ref.^63^), and integrated and adapted a semi-automated iterative procedure (for details, see ref.^65^). For each subject, a subject-specific mask of the cerebral aqueduct was created by identifying high-variance voxels in the region. Following this, subject-specific PAG masks were created by dilating the aqueduct mask (2 voxels, i.e., 2.2 mm) and masking voxels in the original subject-specific aqueduct mask, voxels not in a subject-specific gray-matter segmentation (> 50% probability), and any voxels outside the target range [MNI: −22 > y > −42; z > −14]. A custom group PAG-aligned template was created using DARTEL^108^, which first aligned subject-specific PAG masks in approximate MNI space, then warped whole-brain contrast images using the subject-specific transformation to the common PAG-aligned space.

The group-aligned PAG template was subdivided into columnar region of interests (ROIs) (Fig. 2). Voxel x/y/z coordinates were submitted to PCA to identify the PAG longitudinal axis, and a geometric transformation on the remaining PCA components gave the radius and degree of each PAG voxel (relative to the y axis, at 0°). PAG columns were defined by their degree range: dorsomedial (+/−22.5°), left/right dorsolateral (22.5-60°), left/right lateral (60-97.5°), and left/right ventrolateral (97.5-135°), with the ventromedial quarter (i.e., 90°) of the mask excluded from analysis. The ventral most 90° of the PAG was excluded from analysis, given that it contains other subcortical structures (e.g., dorsal raphe). Columnar ROIs were only used for display purposes, and subdivisions of the PAG used for modeling analyses (Fig. 4b) are described in *Linear Mixed-Effects Model*.

### Peripheral Physiological Recording and Quality Assessment

All peripheral physiological measures were collected at 1kHz using an AD Instruments PowerLab data acquisition system with MR-compatible sensors and LabChart software. Data were continuously acquired throughout the entire scan session and partitioned for alignment with fMRI data using experimenter annotations in LabChart and scanner TR events. A piezo-electric pulse transducer (PowerLab, AD Instruments, Colorado Springs, Colorado, USA) measured heart rate from the left index fingertip. A respiratory belt with a piezo-electric transducer (UFI) measured respiratory rate and was placed around the chest at the lower end of the sternum.

Physiological time-series data were visually inspected for quality in Biopac’s AcqKnowledge data acquisition and analysis software (Biopac Systems Inc., Goleta, California, USA) and in custom visualizations using R (R version 3.6.2^109^) and the ggplot2 package (version 3.2.1^110^). Time-series were classified as high-quality (i.e., few or no motion artifacts), noisy (i.e., IBI: motion artifacts affected < 50% of each 1-minute segment; RR: large, frequent motion artifacts or RRs outside of a typical range of 7-20 breaths per min), and unusable (i.e., IBI: containing excessive artifacts with few reliably detectable peaks; RR: containing low amplitudes that could not be distinguished from noise). Among the 87 participants with usable fMRI data, cardiac data was high-quality (i.e., all 1-minute segments were sufficiently free of motion to easily identify peaks in the IBI signal) for 59 participants, noisy for 12, unusable for 14, and missing for two. Among the 87 participants with usable fMRI data, respiratory data was high-quality for 47 participants, noisy for 20, unusable for 14, and missing for six. Only high-quality data were used for cardiac (IBI) and respiratory (RR) analyses.

### Peripheral Physiological Preprocessing and Analysis

Physiological time-series data were analyzed using MATLAB toolboxes and custom R scripts. Heart rate was calculated using the PhysIO MATLAB toolbox^111^, which used a 0.3 to 9 Hz band-pass filter, and identified peaks using a two-pass process. The first-pass estimated run average heart rate (inverse of IBI), detecting peaks exceeding 40% of normalized amplitude, assuming a minimum peak spacing < 90 beats per minute. First-pass peaks were used to create an averaged template, and the first-pass estimated average heart rate was used as a prior to detect peaks on the second-pass (for more details, see ref.^112^). PhysIO pipeline-detected peaks were compared with peaks identified in Biopac by trained coders. Discrepancies between the two methods were rare (0.5% of peaks across all runs and participants), and occurred very rarely in the dataset scored by a more experienced coder (0.323% of peaks), compared to the dataset scored by a less experienced coder (0.878% of peaks). To reduce drift in the cardiac signal, heart rates were smoothed with a 6-second rolling average and down-sampled into scan repetition time. Upon completion of pre-processing, we then converted heart rate into IBI, as IBI displays a more linear relationship with underlying autonomic cardiac inputs than heart rate^113^. After converting HR to IBI, cardiac data were analyzed exclusively using IBI. Respiratory rates were calculated using custom R scripts. A 1 Hz low-pass filter was applied to the respiratory time-series, and local peaks were identified in a sliding 500 ms window. Small excursions (those < 0.5 SD of average respiratory belt tension across the entire run) were deemed too small to meet criteria as a respiratory peak or trough. To maintain consistency between pre-processing of IBI and RR, RRs were also smoothed with a 6-second rolling average and down-sampled into scan repetition time. Down-sampling IBI and RR into scan repetition time resulted in a total of 263 captured instances of physiological activity for each variable (IBI, RR) in every participant; averages were then computed for each participant and for each resting fixation, 1-back and 3-back task epoch (see *Experimental Paradigm*). Using the averaged task epoch values, change scores were computed as the difference in physiological activity (IBI, RR) between resting fixation and 1-back and 3-back task epoch.

### MRI Analysis

To estimate BOLD signal intensity during the working memory *N*-back task, a general linear model was applied to preprocessed functional time-series (GLM; FSL version 5.0.9). First-level models were estimated for each participant, including separate regressors for the 1-back and 3-back conditions of each trial (1 s). All regressors were convolved with a double gamma hemodynamic response function (HRF). Regressors were modeled in relation to the implicit baseline of the fixation cross. Nuisance regressors included a run intercept, motion (i.e., translation/rotation in x/y/z planes, their mean-centered squares, and their temporal derivatives), aCompCor components (5 cerebral spinal fluid, 5 white matter^114^), a discrete cosine transformation set with a minimum period of 120 seconds, and spike regressors (> 0.5mm framewise displacement^115^). Group-level whole-brain contrast results were thresholded by false discovery rate (*q*_FDR_ < 0.05), with a minimum cluster extent of 100 voxels.

### Linear Mixed-Effects Model

The spatial distribution of PAG BOLD signal intensity increases was analyzed with a linear mixed-effects model, including effects of condition (1-back/3-back), rostral–caudal position rank, PAG degree rank, all interactions, by-participant random slopes, and all by-participant random intercepts. For analysis, PAG BOLD signal intensity was averaged within PAG degree and rostral–caudal position ranks. We used four ordinal, rostral–caudal position ranks for each 3.5 mm segment along the 14 mm rostral–caudal positional axis. We used ten ordinal, dorsomedial to ventrolateral PAG degree ranks for each 13.5° section of the PAG, averaging across bilateral sections (e.g., rank 0 includes 0-13.5° in both left/right PAG, while rank 9 includes 121.5-135° in both left/right PAG).

Analyses were conducted in R (R version 3.6.2^109^) using the `lmer4` package^116^, with degrees of freedom approximated by the Satterthwaite method, as implemented in the `lmerTest` package^117^. The code was entered as follows:

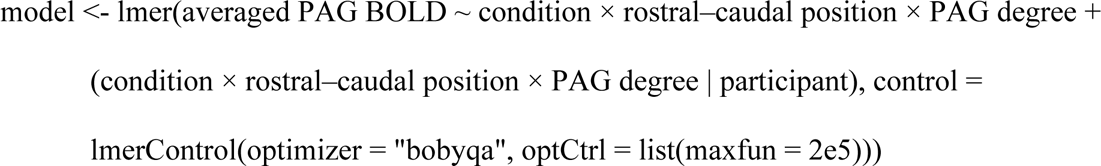

