## Supplementary Materials for "7-Tesla evidence for columnar and rostral–caudal organization of the human periaqueductal gray response in the absence of threat: a working memory study"

Affiliations:

| PAG Afferent Connections | | | | | |
|---|---|---|---|---|---|
| **Region** | **Sub-region** | **DM** | **DL** | **L** | **VL** |
| Amygdala | Anterior amygdaloid nuclei | | | | Rabbit[3] |
| Amygdala | Central amygdaloid nuclei | Cat[1], Monkey[2], Rabbit[3] | Cat[1], Rabbit[3] | Monkey[2], Rabbit[3], Rat[4] | Cat[1], Monkey[2], Rabbit[3] |
| Amygdala | Medial amygdaloids nuclei | Rabbit[3] | Rabbit[3] | Rabbit[3] | Rabbit[3] |
| Amygdala | Ventrolateral basal nucleus of the amygdala | | Monkey[2] | Monkey[2] | |
| Basal Ganglia | Globus pallidus | | | | Rabbit[3] |
| Cerebellum | Dentate nucleus | Rabbit[3] | Rabbit[3] | Rabbit[3] | Rabbit[3] |
| Cerebellum | Fastigial nucleus | | | | Mouse[5], Rabbit[3] |
| Cerebellum | Interpositus nucleus | | | | Rabbit[3] |
| Cingulate | Anterior cingulate | | | | Rat[8] |
| Cingulate | Anterior cingulate area 24 | Monkey[2] | | Monkey[2] | Monkey[2], Rat[6] |
| Cingulate | Dorsal anterior cingulate cortex | | Rat[7] | Rat[7] | Rat[7] |
| Cingulate | Intermediate anterior cingulate | Rat[9] | Rat[9] | | Rat[9] |
| Cingulate | Ventral anterior cingulate cortex | | Rat[7] | | Rat[7] |
| Frontal Lobe | Primary motor cortex | | | | Rat[8] |
| Hippocampus | Hippocampus | *Not present* | | | |
| Hypothalamus | Anterior hypothalamic nuclei | | | | Rat[8] |
| Hypothalamus | Anterior hypothalamus | | | Rat[10] | |
| Hypothalamus | Dorsal premammillary nucleus | Rabbit[3], Rat[11] | Rabbit[3], Rat[11,12] | Rabbit[3,12] | Rabbit[3], Rat[11,8] |
| Hypothalamus | Dorsomedial hypothalamus | Rabbit[3], Rat[13] | Rabbit[3], Rat[13,12] | Rabbit[3], Rat[13,12] | Rat[13] |
| Hypothalamus | Lateral hypothalamus | | | Rat[12] | Rabbit[3], Rat[8] |
| Hypothalamus | Medial preoptic nuclei | Rabbit[3], Rat[4] | Rabbit[3] | Rabbit[3], Rat[4] | Rat[48] |
| Hypothalamus | Parasubthalamic nuclei | | | | Rat[8] |
| Hypothalamus | Periventricular nucleus | | | | Rabbit[3] |
| Hypothalamus | Preoptic nuclei | | | | Rabbit[3] |
| Hypothalamus | Retrochiasmatic area | | | | Rat[8] |
| Hypothalamus | Supramammillary nucleus | | | Rat[12] | Rabbit[3], Rat[12] |
| Hypothalamus | Supraoptic nucleus | | | Rabbit[3] | Rabbit[3] |
| Hypothalamus | Ventromedial nucleus | | | | Rat[8] |
| Insular Cortex | Dorsal insular cortex | | Rat[7] | Rat[7] | Rat[7] |
| Insular Cortex | Insular cortex | | | | Rat[6] |
| Insular Cortex | Posterior insular cortex | | | | Rat[78] |
| Insular Cortex | Rostral insular cortex | Rabbit[3] | Rabbit[3] | Rabbit[3] | |
| Lateral Sulcus | Insular cortex | | | | Rat[6] |
| Locus Coeruleus | Locus coeruleus | | | | Monkey[23] |
| Medial Frontal Cortex | Supplementary eye field | Monkey[14] | Monkey[14] | | Monkey[14] |
| Medial Olfactory Area | Lateral septum | Monkey[15] | Monkey[15] | | |
| Medial Olfactory Area | Ventral septum | Monkey[15] | Monkey[15] | | |
| Medial Prefrontal Cortex | Dorsomedial prefrontal cortex | | Monkey[21] | | |

| | | | | | |
|---|---|---|---|---|---|
| Medial Prefrontal Cortex | Dorsomedial prefrontal cortex area 24 | Rabbit[3] | Rabbit[3] | Rabbit[3] | Rabbit[3] |
| Medial Prefrontal Cortex | Dorsomedial prefrontal cortex area 24b | | | Monkey[2] | |
| Medial Prefrontal Cortex | Dorsomedial prefrontal cortex area 9 | | | Monkey[2] | |
| Medial Prefrontal Cortex | Medial prefrontal area 25 | | Monkey[2,16,17] | | Monkey[2] |
| Medial Prefrontal Cortex | Medial prefrontal area 32 | Monkey[2], Rabbit[3] | Monkey[2,16], Rabbit[3] | Rabbit[3] | Rabbit[3], Rat[6] |
| Medial Prefrontal Cortex | Medial prefrontal cortex | Mouse[18] | Monkey[19], Mouse[18], Rat[20] | Mouse[18] | Rat[20] |
| Medulla Oblongata | Lateral cervical nucleus | | | | Rabbit[3] |
| Medulla Oblongata | Nucleus paragigantocellularis | | | | Rabbit[3] |
| Medulla Oblongata | Raphe obscurus nucleus | | | | Rabbit[3] |
| Medulla Oblongata | Solitary nucleus | | | | Rabbit[3], Rat[22] |
| Medulla Oblongata | Spinal trigeminal nucleus | | | | Rabbit[3] |
| Olfactory Bulb | Olfactory bulb | | | | Rat[6] |
| Olfactory Cortex | Anterior olfactory cortex | | | | Rat[6] |
| Orbital Cortex | Dorsolateral orbital cortex | | Rat[7] | Rat[7] | Rat[7] |
| Orbital Cortex | Medial orbital cortex | | | | Rat[7] |
| Orbital Cortex | Orbital area 12o | | Monkey[2] | | Monkey[2] |
| Orbital Cortex | Orbital area 13a | | Monkey[2] | | Monkey[2] |
| Orbital Cortex | Orbital cortex | | | | Rat[6] |
| Orbital Cortex | Ventral orbital cortex | | | | Rat[7] |
| Orbital Cortex | Ventrolateral orbital cortex | | | | Rat[7] |
| Pons | Caudal pontine reticular nuclei | Rabbit[3] | Rabbit[3] | Rabbit[3] | Rabbit[3] |
| Pons | Pontine reticular formation | Rabbit[3] | Rabbit[3] | Rabbit[3] | Rabbit[3] |
| Pons | Raphe magnus | | | | Rabbit[3] |
| Prefrontal Cortex | Arcuate frontal eye field | Monkey[14] | Monkey[14] | Monkey[14] | Monkey[14] |
| Reticular Formation | Cuneiform nucleus | | | Rat[10] | |
| Reticular Formation | Reticular formation | Cat[1] | Cat[1] | Monkey[24] | Cat[1] |
| Spinal Cord | Dorsal horn | | | | Rabbit[3] |
| Spinal Cord | Spinal cord | Cat[25] | Cat[25] | Cat[25] | Cat[25] |
| Superior Colliculus | Medial superior colliculus | Mouse[29] | Mouse[29] | | |
| Superior Colliculus | Superior colliculus | Cat[26], Monkey[27] | Cat[26], Monkey[27] | Cat[26], Monkey[27] | Monkey[27,28] |
| Thalamus | Parafascicular nucleus | | | | Rabbit [1] |
| Thalamus | Sub-thalamic nucleus | | Monkey[30] | | Rabbit [1] |
| Thalamus | Suprageniculate nucleus | Rabbit[1] | Rabbit[1] | Rabbit[1] | Rabbit[1] |
| Zona Incerta | Zona incerta | Rabbit[1] | Rabbit[1] | Rabbit[1] | Rabbit[1], Rat[6] |

| **PAG Efferent Connections** | | | | | |
|---|---|---|---|---|---|
| **Region** | **Sub-region** | **DM** | **DL** | **L** | **VL** |
| Amygdala | Amygdala | | | | Rabbit[31] |
| Amygdala | Central extended amygdala | | | Rat[32] | Rat[32] |
| Amygdala | Central nucleus of amygdala | | Rat[4] | Rat[4] | Rat[4] |
| Basal Forebrain | Substantia innominata | | | | Rabbit[31] |
| Brainstem | Nucleus incertus | | | | Rat[33] |
| Brainstem | Subcoeruleus nucleus | | Rat[34] | | |
| Hippocampus | Hippocampus | *Not present* | | | |
| Hypothalamus | Posterior hypothalamic area | | | Rat[35] | Rat[35] |
| Hypothalamus | Ventrobasal hypothalamus | Rat[36] | | Rat[36] | Rat[36] |
| Inferior Colliculus | Inferior colliculus | | | Rabbit[31] | Rabbit[31] |
| Locus Coeruleus | Locus coeruleus | Monkey[37] | Monkey[37], Rat[34] | Monkey[37], Rat[38] | Monkey[37], Rabbit [31], Rat[38,39] |
| Medulla Oblongata | Caudal ventrolateral medulla | | | | Rat[40] |
| Medulla Oblongata | Lateral medulla | | | Monkey[41] | Monkey[41] |
| Medulla Oblongata | Lateral paragigantocellularis nucleus | | | | Rat[34] |
| Medulla Oblongata | Nucleus raphe magnus | Cat[42], Monkey[37] | Monkey[37] | Cat[42], Monkey[37] | Cat[42], Monkey[37], Rat[34] |
| Medulla Oblongata | Paragigantocellularis nucleus | | Rat[34] | | Monkey[37] |
| Medulla Oblongata | Rostral ventral medulla | | | | Rat[43] |
| Medulla Oblongata | Rostral ventrolateral medulla | Cat[44], Rat[45] | | Cat[44] | Cat[44], Rat[45] |
| Medulla Oblongata | Rostro-ventromedial medulla | Rat[36] | | Rat[36] | Rat[36] |
| Medulla Oblongata | Solitary nucleus | | Rat[34] | | Rat[34] |
| Medulla Oblongata | Spinal trigeminal nucleus | | | | Monkey[37] |
| Medulla Oblongata | Trigeminal sensory complex | Rat[46] | Rat[46] | Rat[46] | Rat[46] |
| Pons | Lateral parabrachial nucleus | Rat[47] | | Rat[47] | Rat[47] |
| Pons | Medial parabrachial nucleus | Rat[47] | | Rat[47] | Rat[47] |
| Pons | Parabrachial nucleus | Rat[47] | Rat[47] | Rat[47] | Rat[47] |
| Pons | Reticulotegmental nucleus | | | | Rabbit[31] |
| Pons | Ventrolateral pons | Monkey[48] | Monkey[48] | Monkey[48] | Monkey[48] |
| Reticular Formation | Caudal ventrolateral reticular nucleus | | | | Rat[34] |
| Reticular Formation | Gigantocellular reticular nucleus | | Rat[34] | | |
| Reticular Formation | Nucleus ambiguus | | Rat[34] | Monkey[4937] | Rat[34] |
| Reticular Formation | Nucleus cuneiformis | Monkey[37], Rabbit[31] | Monkey[37], Rabbit[31] | Rabbit[31] | Monkey[37], Rabbit[31] |
| Reticular Formation | Reticularis magnocellularis | Cat[42] | | Cat[42] | Cat[42], Monkey[37] |
| Reticular Formation | Reticularis pontis oralis | Monkey[37] | Monkey[37] | Monkey[37] | Monkey[37] |
| Reticular Formation | Retroambiguus nucleus | | | Monkey[4150] | Monkey[50] |
| Reticular Formation | Rostral ventrolateral reticular nucleus | | | | Rat[34] |
| Spinal Cord | Spinal cord | | | Monkey[51] | Monkey[51] |
| Substantia Nigra | Substantia nigra | | | | Rat[43] |
| Superior Colliculus | Superior colliculus | Rabbit[31] | Rabbit[31] | | Rabbit[31], Rat[39] |

| | | | | | |
|---|---|---|---|---|---|
| Thalamus | Caudal paraventricular nucleus | Rat[52] | Rat[52] | Rat[52] | Rat[52] |
| Thalamus | Central lateral thalamic nucleus | Rat[53] | Rat[53] | Rat[53] | Rat[53] |
| Thalamus | Central medial thalamic nucleus | Rat[53] | | Rat[53] | Rat[53] |
| Thalamus | Intermediate paraventricular nucleus | Rat[52] | Rat[52] | Rat[52] | Rat[52] |
| Thalamus | Lateral parafascicular thalamic nucleus | Rat[53] | | Rat[53] | |
| Thalamus | Medial parafascicular thalamic nucleus | | | Rat[53] | Rat[53] |
| Thalamus | Medial thalamus | Rat[36] | | Rat[36] | Rat[36] |
| Thalamus | Mediodorsal thalamic nucleus | | Rat[53] | Rat[53] | Rat[53] |
| Thalamus | Paracentral thalamic nucleus | | Rat[53] | Rat[53] | Rat[53] |
| Thalamus | Parafascicular thalamic nucleus | | | | Rat[39] |
| Thalamus | Paratenial thalamic nucleus | | | Rat[53] | |
| Thalamus | Rostral paraventricular nucleus | Rat[52] | Rat[52] | Rat[52] | Rat[52] |
| Thalamus | Sub-thalamic nucleus | | | | Rabbit[31] |
| Thalamus | Submedius thalamic nucleus | Rat[53] | Rat[53] | Rat[53] | Rat[53] |
| Thalamus | Subparafasicular thalamic nucleus | | Rat[53] | | |
| Thalamus | Suprageniculate nucleus of the posterior thalamus | | | | Rabbit[31] |
| Thalamus | Ventromedial thalamic nucleus | | | Rat[53] | Rat[53] |
| Ventral Midline of the Thalamus | Reuniens thalamic nucleus | | Rat[53] | Rat[53] | Rat[53] |
| Ventral Midline of the Thalamus | Rhomboid thalamic nucleus | | | Rat | Rat[53] |
| Ventral Tegmental Area | Ventral tegmental area | | | Mouse[54] | Mouse[54], Rat[43] |
| Zona Incerta | Zona incerta | Rat[53] | Rat[53] | Rat[53] | Rat[39] |

**Supplementary Table 1.** Monosynaptic afferent and efferent connections of the PAG. Species in which connections were observed are listed in cells, with relevant citations listed below. Blank cells denotes the *absence of evidence* for or against a connection, to the best of our knowledge. Cells labeled "not present" in gray text denotes a study finding *evidence of absence.* In order for a monosynaptic connection to be included in the table, monosynaptic connections needed to exhibit PAG columnar specificity. The PAG receives afferent (top-down) and efferent (bottom-up) connections from both cortical and subcortical areas of the brain. Projections arising from cortical regions include the prefrontal cortex (An et al., 1998; Floyd et al., 2000; Hardy & Leichnetz, 1981), insular cortex (Benarroch, 2012; Neafsey et al., 1986), and the motor cortex

(Beitz, 1982; Linnman et al., 2012; Mantyh, 1983a). Projections arising from subcortical and hindbrain areas, include the hypothalamus (Behbehani et al., 1988), amygdala (Rizvi et al., 1991) brainstem (Beitz et al., 1986), and spinal cord (Behbehani, 1995). The PAG's efferents project to brain areas that directly modulate the motor, behavioral, and sympathetic responses that operate to meet the demands reported by ascending sensory information. These brain areas include the thalamus (Krout & Loewy, 2000), midbrain (Meller & Dennis, 1991), and brainstem (Mantyh, 1983b).

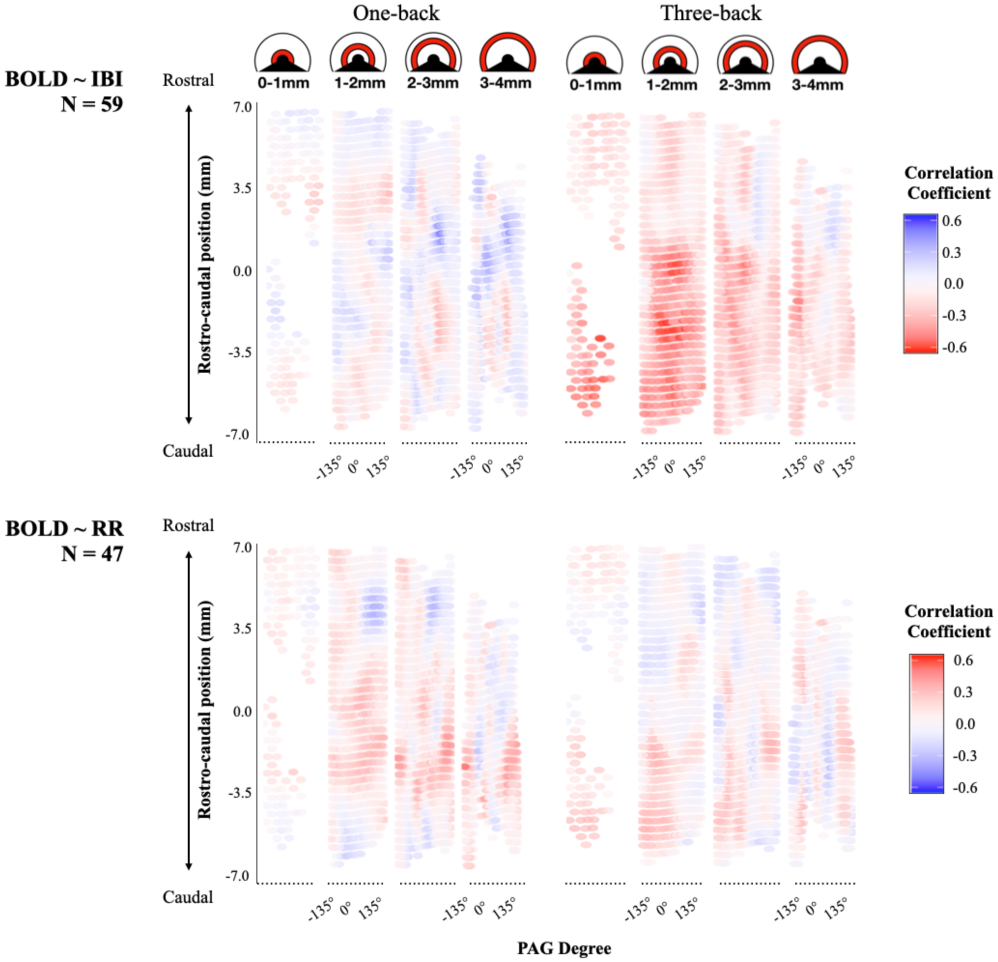


**Supplementary Figure 1.** Between-subject correlations between peripheral physiology voxelwise PAG BOLD signal intensity.

Upper panels display radial cross-sections of the PAG cylinder (e.g., 0-1 mm). The y-axis illustrates voxel position along the rostral–caudal axis. The x-axis represents voxel position in +/− 135 degrees from the brain midline (ventrolateral = +/− 97.5-135°; lateral = +/− 60-97.5°; dorsolateral = +/− 22.5-60°; dorsomedial = +/− 22.5°). The heat map illustrates magnitude (i.e., strength) of the correlation coefficients. Abbreviations: IBI: interbeat interval; RR: respiration rate.
